# Prototyping 1,4-butanediol (BDO) biosynthesis pathway in a cell-free transcription-translation (TX-TL) system

**DOI:** 10.1101/017814

**Authors:** Yong Y. Wu, Stephanie Culler, Julia Khandurina, Stephen Van Dien, Richard M. Murray

## Abstract

Current methods for assembling metabolic pathways require a process of repeated trial and error and have a long design-build-test cycle. Further, it remains a challenge to precisely tune enzyme expression levels for maximizing target metabolite production. Recently it was shown that a cell-free transcriptional-translation system (TX-TL) can be used to rapidly prototype novel complex biocircuits as well as metabolic pathways. TX-TL systems allow protein expression from multiple DNA pieces, opening up the possibility of modulating concentrations of DNA encoding individual pathway enzymes and testing the related effect on metabolite production. In this work, we demonstrate TX-TL as a platform for exploring the design space of metabolic pathways using a 1,4-BDO biosynthesis pathway as an example. Using TX-TL, we verified enzyme expression and enzyme activity and identified the conversion of 4-hydroxybutyrate to downstream metabolites as a limiting step of the 1,4-BDO pathway. We further tested combinations of various enzyme expression levels and found increasing downstream enzyme expression levels improved 1,4-BDO production.

## Introduction

Designing and testing biosynthesis pathways *in vivo* is a time-consuming and labor-intensive process. Cloning a pathway into plasmids *in vivo* has a one-week testing cycle [1]. For example, it took 150 person-years using this method to develop and implement production process of the anti-malaria drug artemisinin [2, 3]. On the other hand, cell-free transcription-translation (TX-TL) systems can be utilized to reduce cloning iterations and streamline the process of verifying enzyme expression. The design-build-test cycle of a biological circuit using linear DNAs in TX-TL takes less than one day [4], which can dramatically reduce research efforts. TX-TL allows simultaneous protein expression from multiple pieces of DNA, including linear DNA. Such properties can help quickly verify pathway enzyme expression and test pathway enzyme activity before a pathway is integrated *in vivo*.

Metabolic pathways consisting of multiple parts and factors should be combined in precise combinations to achieve desired functions. Enzyme expression levels and enzyme activity affect target product yield. However, tuning enzyme expression requires engineering on the level of transcription, translation, and enzyme activity. Further, balancing expression of multiple genes in parallel remains a challenge. Currently, enzymes are overexpressed when they are identified to be important for improving target metabolite production. Protein overexpression can affect cell growth and possibly reduce metabolite production. A few groups have previously demonstrated the feasibility of tuning protein expression levels *in vivo* for improve metabolite productions [5-7]. This work aims to utilize TX-TL to rapidly tune pathway enzyme expression *in vitro* for improved metabolite productions.

A 1,4-butanediol (BDO) biosynthesis pathway has been prototyped by 1) verifying enzyme expression and activity in TX-TL, 2) modulating enzyme expression levels in TX-TL for improved metabolite production, and 3) reconstructing the pathway *in vivo*. Overall workflow is shown in Figure 1. 1,4-BDO is an important commodity chemical with an estimated global market of 2.5 million ton by 2017[8]. Genomatica Inc. (San Diego, CA) previously has reported the production of 18g/L 1,4-BDO using an engineering *E. coli* strain [9]. Pathway enzymes include succinyl-CoA synthetase (*sucCD*), CoA-dependent succinate semialdehyde dehydrogenase (*sucD*), 4-hydroxybutyrate dehydrogenase (*4-hbd*), 4-hydroxybutyryl-CoA transferase (*cat2*), 4-hydroxybutyryl-CoA reductase (*ald*) and alcohol dehydrogenase (*adh*). The 1,4-BDO pathway is shown below in Figure 2.

**Figure 1:**
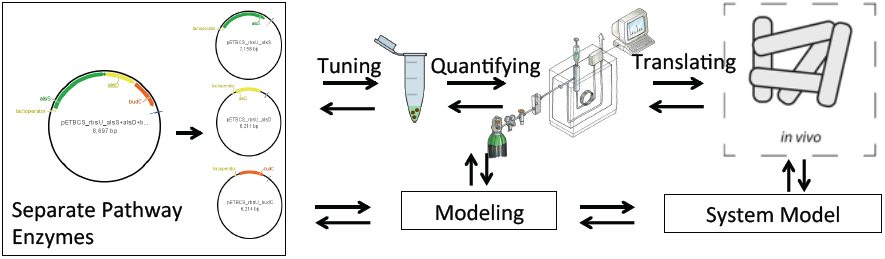
Cell-free biomolecular breadboard for pathway design space exploration. From left to right, cloning gene sequence of pathway enzymes onto individual vectors and under the same promoter for verifying enzyme expression and activity, modulating enzyme expression levels for improved metabolite production in cell-free TX-TL systems, correlating enzyme expression in TX-TL to *in vivo*, and eventually optimizing pathway design *in vivo*. In parallel, mathematical models can be developed to identify key parameter for understanding pathway dynamics.

**Figure 2:**
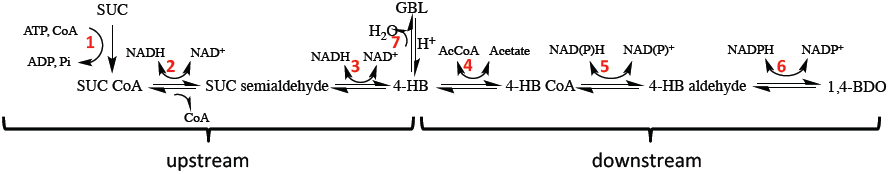
1,4-BDO biosynthesis pathway introduced to TX-TL: Enzymes for each numbered reactions are: 1) succinyl-CoA synthetase; 2) CoA-dependent succinate semialdehyde dehydrogenase; 3) 4-hydroxybutyrate dehydrogenase; 4) 4-hydroxybutyryl-CoA transferase; 5) 4-hydroxybutyryl-CoA reductase; 6) alcohol dehydrogenase. Reaction 7 is a spontaneous reaction of 4-HB converting to *gamma*-butyrolactone (GBL). Reactions 1, 2, and 3 are considered upstream reactions (abbreviated as U), and reactions 4, 5, and 6 are considered downstream reactions (abbreviated as D).

## Result and Discussion

Pathway enzymes were expressed in TX-TL reactions by adding linear DNA. Linear pieces of DNA were generated by amplifying regions encoding a promoter, a 5’-untranslated region (UTR), a coding sequence of an individual pathway enzyme, and a terminator. End products of TX-TL reactions were directly used for polyacrylamide gel electrophoresis with sodium dodecyl sulfate (SDS-PAGE) preparation, and all enzymes of the 1,4-BDO pathway showed up on the gel at expected sizes (as shown in Figure 3). Enzyme activity was verified by testing downstream enzymes and upstream enzymes separately. The downstream module consists of three enzymes that facilitate reactions after the synthesis of 4-hydroxybutyrate (4-HB), and the upstream module consists of three enzymes that facilitate the rest of the pathway reactions. Downstream and upstream modules are shown in Figure 2. Upstream enzymes were tested by adding 30 mM succinate in TX-TL reaction, while downstream enzymes were tested by adding 30 mM of a surrogate compound butyrate. TX-TL reactions were analyzed using GC-MS for expected metabolites, which was 4-HB and butanol. Both were detected: 4-HB at about 14 mM and butanol at about 1 mM.

**Figure 3:**
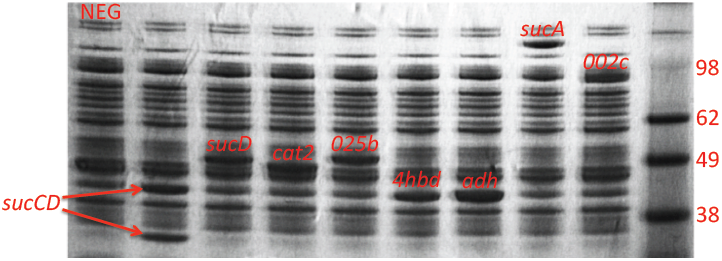
Verifying enzyme expression using SDS-PAGE gel: From left to right are negative control (NEG), succinyl-CoA synthetase (*sucCD*, two subunits *α* and *β* at 29.6 kDa and 41.4 kDa [10]), CoA-dependent succinate semialdehyde dehydrogenase (*sucD*, 50.2 kDa), 4-hydroxybutyryl-CoA transferase (*cat2*, 48.0 kDa), 4-hydroxybutyryl-CoA reductase (*ald-025b*, 52.1 kDa), 4-hydroxybutyrate dehydrogenase (*4-hbd*, 41.3 kDa) alcohol dehydrogenase (*adh*, 43.1 kDa), 2-oxoglutarate decarboxylase (*sucA*, 100kDa), 4-hydroxybutyryl-CoA reductase (*ald-002c*, 95.3kDa), and protein ladder.

Enzyme expression levels were modulated in TX-TL by varying concentrations of linear DNA encoding individual enzymes. Initially, combinations of different concentrations of linear DNA encoding upstream (U) and downstream (D) enzymes were tested. Enzyme *ald-002c* was used. Up to 1 mM of 1,4-BDO was detected from the linear DNA ratio of 4U:8D, which was about 30% more than 1,4-BDO resulted from the ratio of 8U:8D and 100% more than 1,4-BDO resulted from ratio of 4U:4D (shown in Figure 4a). Related metabolites were also measured (shown in Figure 4b). Most of the substrates got converted into 4-HB or *γ*-butyrolactone (GBL), and downstream reactions were suspected to be bottlenecks of the 1,4-BDO pathway.

**Figure 4:**
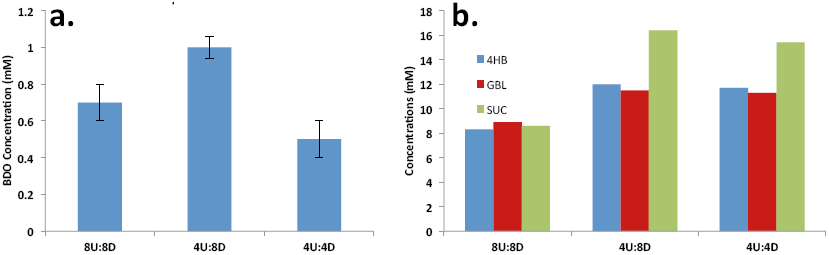
**a.** 1,4-BDO produced in TX-TL reactions after 18 hours: error bars were estimated based on measurements from GC-MS and LC-MS. x-axis notation: “8U:8D” denotes for 8 nM of linear DNA encoding upstream enzymes and 8 nM of linear DNA encoding downstream enzymes. **b.** Related metabolites produced in the same TX-TL reactions as in **a** after 18 hours: 4HB—4-hydroxybutyrate, GBL–*gamma*-butyrolactone, and SUC—succinate.

To further explore the design space of the 1,4-BDO pathway, two different *ald*s were tested: *002c* and *025b*. *ald-002c* is a bi-functional enzyme [9] which catalyzes both reaction 5 and 6 shown in Figure 2. Figure 5a shows that TX-TL reactions added with *ald-002c* generally produced more 1,4-BDO than reactions added with *ald-025b*, regardless of enzyme expression levels. Reactions with *ald-025b* produced around 0.3 mM of 1,4-BDO. TX-TL reactions added with only *ald-002c* produced 1,4-BDO comparable to reactions added with both *ald-002c* and *adh*. Figure 5b again confirms metabolic flux accumulates as 4-HB or GBL and shows significant lactate accumulation. Pyruvate might be converted into lactate as part of the mechanism for regenerating ATP, which is important for protein synthesis [11].

**Figure 5:**
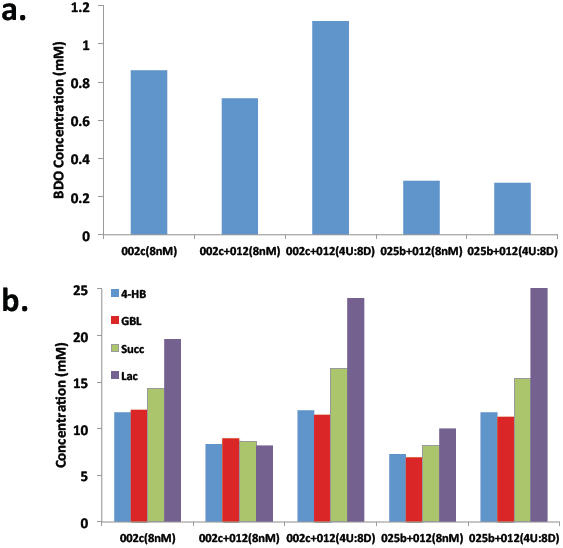
a. 1,4-BDO produced in TX-TL reactions after 18 hours. x-axis notation: “002c (8nM)” denotes for 8 nM of linear DNA encoding upstream enzymes, 4-hydroxybutyryl-CoA transferase (cat2), and *ald-002c*. “002c +012 (8nM)” denotes for 8 nM of linear DNA encoding upstream enzymes, cat2, and *ald-002c* and *adh-012*. Error bars were estimated based on measurements from GC-MS and LC-MS. b. Related metabolites produced in the same TX-TL reactions as in **a** after 18 hours: 4HB (4-hydroxybutyrate), GBL (*γ*-butyrolactone), Succ (succinate), and Lac (lactate).

Data so far has indicated that downstream reactions are pathway bottlenecks. One possibility is that characteristics of downstream enzymes might affect metabolite production, and the reversibility of pathway enzymes were tested. 1,4-BDO and 4-HB were added into TX-TL reactions to test the reversibility of downstream enzymes. Figure 6 shows that the downstream enzymes do not convert significant amount of 1,4-BDO or 4-HB into respective upstream metabolites. However, significant amount of GBL was generated when 4-HB is added. It is likely that the spontaneous conversion from 4-HB to GBL is the key reaction that causes low conversion of 1,4-BDO.

**Figure 6:**
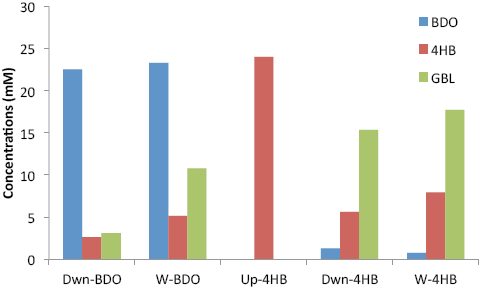
Testing pathway reversibility. x-axis notation: “Dwn-BDO” denotes for 1,4-BDO added to TX-TL reaction with linear DNA encoding downstream enzymes. Dwn—downstream enzymes, W—both downstream and upstream enzymes, and Up—upstream enzymes.

## Conclusion/Future Work

Data above show that using linear DNA in TX-TL can tune pathway enzyme expression for improved metabolite productions. This set of experiments also demonstrates TX-TL’s ability to quickly test pathway reactions of interest without significant cloning or cell transformation. In the future, more concentration combinations of linear DNA encoding individual enzymes will be tested for improved metabolite production. Enzyme expression levels will be tuned using standardized transcription and translation initiation elements. Translational coupling architectures called bicistronic designs (BCDs) will help tune enzyme expression levels more reliably [12]. BCDs will be used for designing metabolic pathways to reflect enzyme expression levels that result with high metabolite production in TX-TL. Future work will also include reconstructing 1,4-BDO pathway *in vivo* using enzyme sequences designed with BCDs. Constructs for individual enzymes will contain a promoter, a BCD, coding sequences of individual enzymes, and a terminator. Individual enzyme constructs will be placed on plasmids with spacing to minimize crosstalk.

## Materials and Methods

### Cell-Free Expression Preparation and Execution

Preparation of the cell-free TX-TL expression system was done according to protocols previously described by colleagues [13], resulting in extract with conditions: 8.9-9.9 mg/mL protein, 4.5-10.5 mM Mg-glutamate, 40-160 mM K-glutamate, 0.33-3.33 mM DTT, 1.5 mM each amino acid except leucine, 1.25 mM leucine, 50 mM HEPES, 1.5 mM ATP and GTP, 0.9 mM CTP and UTP, 0.2 mg/mL tRNA, 0.26 mM CoA, 0.33 mM NAD+, 0.75 mM cAMP, 0.068 mM folinic acid, 1 mM spermidine, 30 mM 3-PGA, 2% PEG-8000. When possible, inducers such as IPTG, pyruvate, and linear DNA were added to a mix of extract and buffer to ensure uniform distribution. TX-TL reactions were conducted in PCR tubes and kept at 29°C with incubation in PCR machine.

### Plasmid DNA and PCR Product Preparation

PCR products were amplified using Pfu Phusion Polymerase (New England Biolabs). Plasmids were miniprepped using a QIAprep spin columns (Qiagen). All plasmids were processsed at stationery phase. Before use in the cell-free reaction, both plasmids and PCR products underwent an additional PCR purification step using a DNA Clean & Concentrator column (Zymo Research), which removed excess salt detrimental to TX-TL, and were eluted and stored in 10 mM Tris-Cl solution, pH 8.5 at 4°C for short-term storage and -20°C for long-term storage.

## Acknowledgement

We thank Genomatica for providing the materials used in this work, including plasmids encoding enzymes for the 1,4-BDO biosynthesis pathway and strain ECKh-422. We thank Nathan Dalleska and the Environmental Analysis Center for the support and assistance using GC/MS. This material is based upon work supported in part by the Defense Advanced Research Projects Agency (DARPA/MTO) Living Foundries program; contract number HR0011-12-C-0065 (DARPA/CMO). Y.Y.W is supported by NIH/NRSA Training Grant 5 T32 GM07616. The views and conclusions contained in this document are those of the authors and should not be interpreted as representing officially policies, either expressly or implied, of the Defense Advanced Research Projects Agency or the U.S. Government.

## Reference

1 Alzari, P.M., et al., Implementation of semi-automated cloning and prokaryotic expression screening: the impact of SPINE. Acta Crystallographica Section D, 2006. 62(10): p. 1103–1113.

2 Kwok, R., Five hard truths for synthetic biology. Nature, 2010. 463: p. 288–290.

3 Ro, D.-K., et al., Production of the antimalarial drug precursor artemisinic acid in engineered yeast. Nature, 2006. 440(7086): p. 940–943.

4 Sun, Z.Z., et al., Linear DNA for Rapid Prototyping of Synthetic Biological Circuits in an Escherichia coli Based TX-TL Cell-Free System. ACS Synthetic Biology, 2013.

5 Du, J., et al., Customized optimization of metabolic pathways by combinatorial transcriptional engineering. Nucleic Acids Research, 2012. 40(18): p. e142.

6 Lee, M.E., et al., Expression-level optimization of a multi-enzyme pathway in the absence of a high-throughput assay. Nucleic Acids Research, 2013.

7 Coussement, P., et al., One step DNA assembly for combinatorial metabolic engineering. Metabolic Engineering, 2014. 23(0): p. 70–77.

8 *1,4-butanediol (BDO) Market By Technology (Reppe Process, Davy process, Butadiene Process, Propylene Oxide Process & Others)*, Applications, & Geography: Global Industry Trends & Forecasts to 2017. 2012 [April 6, 2015]; Available from: http://www.marketsandmarkets.com/Market-Reports/1-4-butanediol-market-685.html.

9 Yim, H., et al., Metabolic engineering of Escherichia coli for direct production of 1,4-butanediol. Nat Chem Biol, 2011. 7(7): p. 445–452.

10 Fraser, M.E., et al., A detailed structural description of Escherichia coli succinly-CoA synthetase. Journal of Molecular Biology, 1999. 288(3): p. 501.

11 Kim, D.-M. and J.R. Swartz, Regeneration of adenosine triphosphate from glycolytic intermediates for cell-free protein synthesis. Biotechnology and Bioengineering, 2001. 74(4): p. 309–316.

12 Mutalik, V.K., et al., Precise and reliable gene expression via standard transcription and translation initiation elements. Nat Meth, 2013. 10(4): p. 354–360.

13 Sun, Z.Z., et al., Protocols for Implementing an Escherichia coli Based TX-TL Cell-Free Expression System for Synthetic Biology. 2013(79): p. e50762.

